# Optimal design of adaptively sampled NMR experiments for measurement of methyl group dynamics with application to a ribosome-nascent chain complex

**DOI:** 10.1101/2020.10.12.336511

**Authors:** Christopher A. Waudby, Charles Burridge, John Christodoulou

## Abstract

NMR measurements of cross-correlated nuclear spin relaxation provide powerful probes of polypeptide dynamics and rotational diffusion, free from contributions due to chemical exchange or interactions with external spins. Here, we report on the development of a sensitivity-optimized pulse sequence for the measurement of cross-correlated relaxation in methyl spin systems by analysis of the differential relaxation of transitions within the ^13^C multiplet. We describe the application of optimal design theory to implement a real-time ‘on-the-fly’ adaptive sampling scheme that maximizes the accuracy of the measured rate constants. The increase in sensitivity obtained using this approach enables, for the first time, quantitative measurements of rotational diffusion within folded states of translationally-arrested ribosome–nascent chain complexes of the FLN5 filamin domain, and can be used to place strong limits on interactions between the domain and the ribosome surface.

## Introduction

Protein biosynthesis by the ribosome is a process of fundamental importance to the cell, ensuring that nascent polypeptides attain their functional native states while avoiding misfolding or aggregation (*1*). Elaborate quality control networks have evolved to chaperone folding proteins, and to sense and eliminate misfolded states (*2*), but for many proteins a key additional facet of efficient biosynthesis is their progressive, ‘co-translational’ folding while still attached to the parent ribosome (*3*). Co-translational folding can reduce the effective volume of conformational space accessible to a nascent polypeptide, facilitating the search for the native state, but it is increasingly apparent that this process can also be modulated or regulated by the ribosome itself (*4*). In particular, a variety of polypeptides have now been observed or inferred to interact with the ribosome surface during translation. This has been associated with perturbations to the free-energy surfaces for folding, in the majority of cases leading to a destabilisation of the nascent polypeptide chain (NC) (*5–10*).

Despite the importance of understanding their influence on co-translational folding processes, at a technical level the direct quantification of interactions between ribosomes and nascent polypeptide chains (NCs) is challenging. Such interactions are intramolecular equilibria, which cannot be characterised via titrations as is common for intermolecular interactions. The NC must also be resolved against typically large background signals from the 2.4 MDa (bacterial) 70S ribosome particle (the large size of which also limits its concentration to a maximum of ca. 10 uM, or 24 mg/mL). One fruitful approach has been the incorporation of fluorescent probes into NCs using non-natural amino acids, for the measurement of time-resolved fluorescence depolarisation (*7*). This technique was used to identify large (ca. 60–90%) populations of a slowly tumbling species, likely ribosome bound, in NCs of the intrinsically disordered protein PIR. However, an intrinsic limitation of this approach is that motions on timescales longer than the fluorophore lifetime (a few ns) cannot be observed.

Solution-state NMR spectroscopy provides an alternative experimental approach to probe co-translational folding (*11*). Despite the large size of the ribosome (with a rotational correlation time of 2.5 μs at 298 K (*12*)), NCs have been found to have sufficient independent mobility to be observable by solution NMR methods (*13*). Translationally-arrested ribosome–nascent chain complexes (RNCs) can be prepared either using cell-free *in vitro* translation systems (*13*, *14*) or exploiting naturally occurring arrest peptides such as SecM (*15*) to prepare and purify RNCs *in vivo* (*8*, *16*, *17*). The majority of ribosomal resonances are effectively suppressed by extremely rapid transverse relaxation associated with their slow tumbling, leading to spectra remarkably free from background. A small number of sharp resonances remain, associated with the mobile L12 stalk protein (*18–20*), but these may be reduced for *in vivo* samples using pulsed isotopic labelling strategies (*17*).

In our prior work, we have studied extensively the co-translational folding of the FLN5 immunoglobulin domain from the multidomain filamin protein of *Dictyostelium discoideum*, using a library of constructs comprising the FLN5 domain attached to the ribosome via varying lengths of the subsequent FLN6 domain (Fig. 1A,B) (*9*, *13*, *14*, *16*, *21*). We have observed site-specific line broadening in unfolded NCs, which are likely to reflect interactions with the ribosome surface, and we have speculated that these interactions may partly underlie the observed delay in the initiation of co-translational folding several residues beyond the point at which the domain is fully emerged from the ribosome exit tunnel (*9*, *21*). However, much less is known about the dynamics of folded FLN5 NCs, interactions of which would perturb the other side of the folding equilibrium. Therefore, in the remainder of this work we will focus exclusively on such systems.

**Figure 1.**
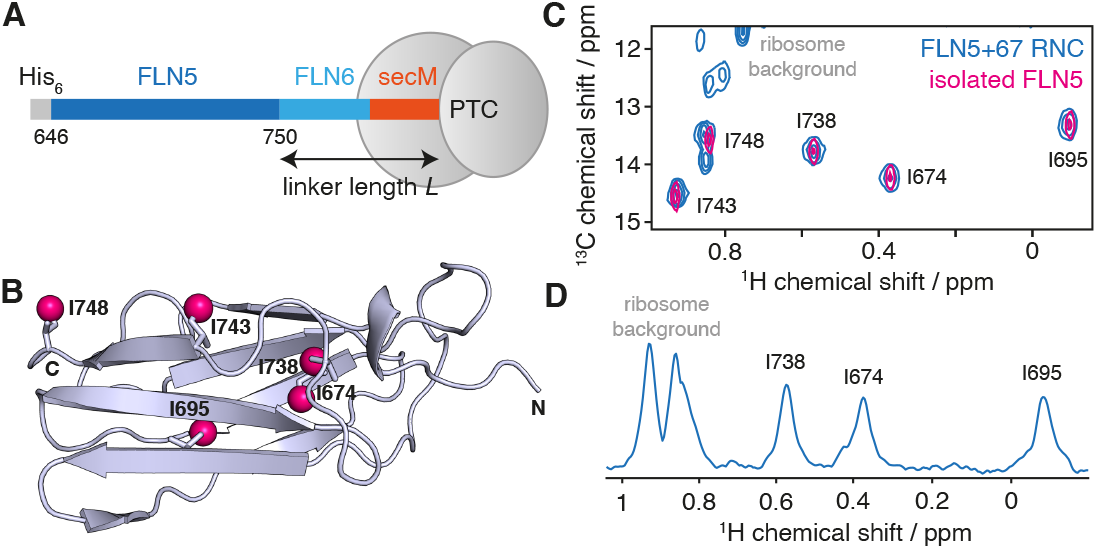
Solution-state NMR spectroscopy of FLN5+6 RNCs. (**A**) Design of RNC constructs (*9*), showing the complete FLN5 filamin domain (646–750) flanked by an N-terminal His_6_ affinity tag, and the subsequent FLN6 domain and 17 amino acid secM arrest motif, of total length *L*. (**B**) Crystal structure (1qfh (*22*)) of FLN5 showing the location of the five isoleucine residues, with Cδ atoms highlighted as magenta spheres. (**C**) ^1^H,^13^C HMQC correlation spectrum of a [^2^H,^13^CH_3_-Ile]-labelled FLN5+67 RNC (blue) and isolated [^2^H,^13^CH_3_-Ile]-labelled FLN5 (298 K, 950 MHz). (**D**) 1D spectrum showing the first increment of an ^1^H,^13^C HMQC measurement of the FLN5+67 RNC.

The first observation of a folded NC by NMR was the FLN5+110 RNC (where 110 refers to the combined length of the FLN6 linker and, if present, arrest peptide), using a uniform ^1^H,^15^N labelling scheme and cell-free expression (*13*). This was followed by the observation of sidechain methyl resonances, in the same system, using a uniform ^1^H,^13^C labelling scheme (*14*). Variations in resonance intensities were observed that were tentatively attributed to altered dynamics within the RNC. However, later measurements, incorporating more stringent controls for NC release, and employing a more sensitive [^2^H,^13^CH_3_-Ile]-labelling scheme (Fig. 1B–D), indicated that previous measurements may have contained additional contributions arising from small amounts of released material (*9*). Large variations in resonance intensity were observed at different linker lengths, indicating that the mobility of the FLN5 domain may be significantly reduced when tethered close to the ribosome surface by a short linker (*9*). To explore these observations further, in the present work we develop an experimental approach suitable for characterising the dynamics within FLN5 RNCs.

We focus here on the analysis of relaxation within isolated methyl ^13^CH_3_ spin systems, which as noted above provide sensitive detection together with good dispersion of chemical shifts. Moreover, methyl spin systems provide a rich array of transitions that may be interrogated to yield dynamic information (Fig. 2) (*23*, *24*). Relaxation rates of these transitions are highly sensitive to restricted rotational diffusion or transient interactions with large species such as the ribosome surface (i.e. transferred relaxation), but also contain contributions from chemical exchange and interactions with external spins. These complicating effects can be eliminated by measurement of cross-correlated relaxation rates, i.e. spin state-dependent differences in relaxation rates, in order to provide a ‘pure’ measurement of rotational diffusion, encoded by the spectral densities *J*(ω). For methyl groups in slowly tumbling macromolecules, the spectral density function is dominated by the zero frequency term, *J*(0) = S^2^τ_c_, where S^2^ is an order parameter for motion of the three-fold symmetry axis, and τ_c_ is the (effective) rotational correlation time (*23*). As S^2^ order parameters are determined by the protein structure, provided that this structure is not perturbed on the ribosome – which would be evident from chemical shift perturbations – changes in *J*(0) in RNCs relative to the isolated protein may then be interpreted simply in terms of changes in effective rotational correlation times.

**Figure 2.**
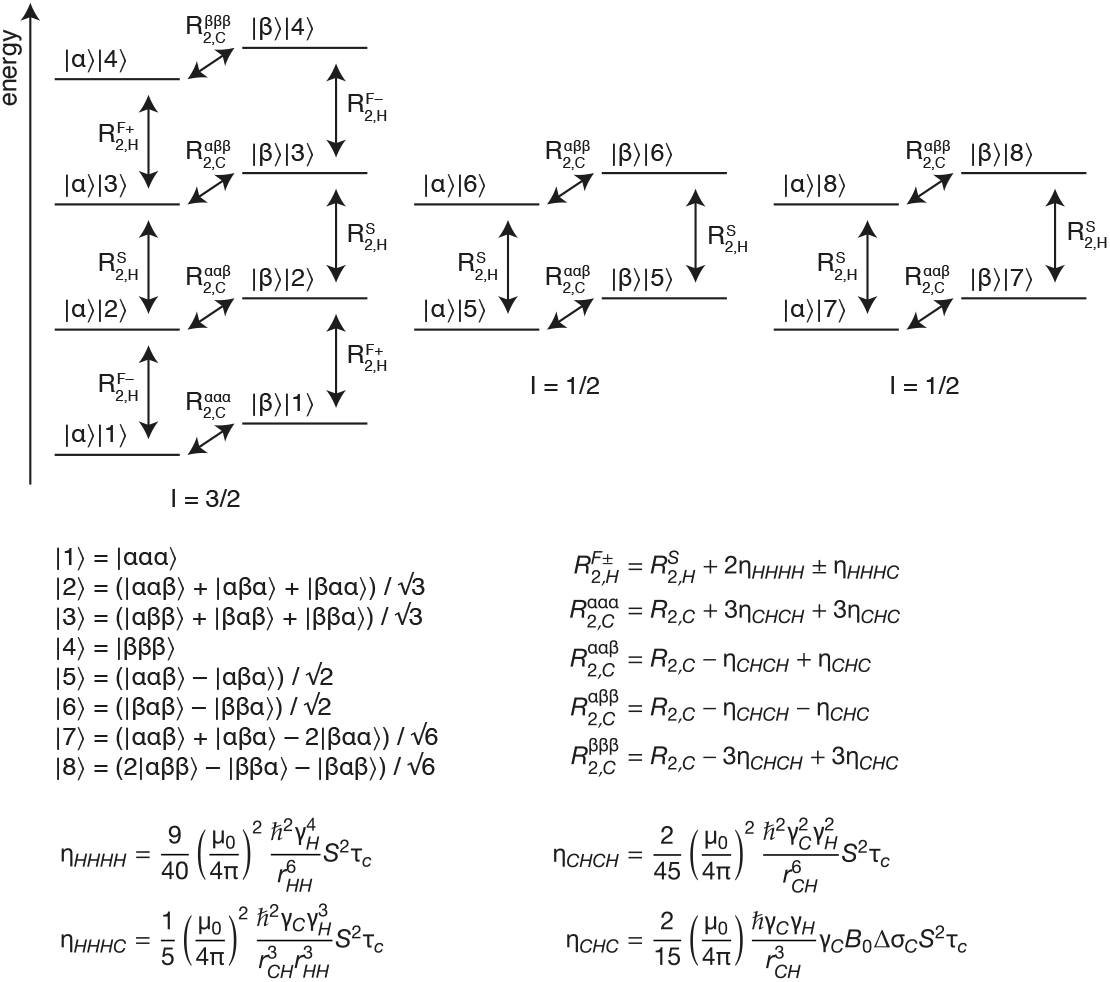
Energy level diagram for a ^13^CH_3_ spin system. Energy levels are labelled as the product of ^13^C and ^1^H spin states, |C〉|H〉, where the numbered ^1^H eigenstates are symmetrised linear combinations of the individual spin states as indicated (*25*). ^1^H eigenstates can be grouped into three independent manifolds and the spin quantum numbers, *I*, for these are shown below the diagram. Single quantum ^1^H and ^13^C transitions are indicated with arrows alongside their relaxation rates, expressions for which are provided below (*26*).

A variety of experiments have been reported for the measurement of rotational diffusion via cross-correlated relaxation in ^13^CH_3_ spin systems (*25–33*). The majority of these fall into two categories. First, measurements of HH/HH dipole-dipole/dipole-dipole (DD/DD) cross-correlated relaxation during the evolution of a ^1^H single quantum coherence, which can be detected either by direct measurement of the slow and fast ^1^H relaxation rates 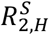 and 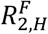 (Fig. 2) (*25*), or via the build-up of double quantum (*27*) or, more sensitively, triple quantum ^1^H coherences (*28*) relative to the relaxation of the single quantum coherence. As ^1^H transitions are degenerate, multiple experiments must be acquired to isolate the various coherences being observed in these approaches. Second, measurements of differential relaxation in the ^13^C multiplet, which is dominated by CH/CH DD/DD cross-correlated relaxation, with additional contributions from CH/C DD/CSA (chemical shift anisotropy) cross-correlations. As ^13^C transitions are not degenerate, this differential relaxation can be read out using a constant-time ^13^C evolution period (*26*), most simply using an F1-coupled constant-time HSQC experiment (*29*, *30*) or, as applied to isolated lysine ^15^NH_3_ spin systems, using a standard F1-coupled HSQC (*34*).

Due to the low concentration of ribosome samples, together with the limited stability of attached nascent chains, optimisation of experimental sensitivity is a critical aspect of RNC NMR, and to this end we have previously explored the use of paramagnetic longitudinal relaxation enhancement agents (*35*) and non-uniform weighted sampling strategies (*10*, *36*). In this regard, the potential to determine cross-correlated relaxation rates from the analysis of multiplet structure in a single, sensitive 2D spectrum is highly attractive. In particular, in this work we focus on measurements of isolated ^13^CH_3_ spin systems against a perdeuterated background, using a conventional (non-constant time) F1-coupled HSQC experiment in order to minimize unnecessary relaxation delays and so maximise sensitivity (*34*). However, careful analysis of this sequence is required in order to account fully for CCR processes during all stages of the pulse sequence, some of which have been overlooked in previous works (*26*, *29*, *30*, *34*). Therefore, here we present a complete expression for the evolution of magnetisation during this experiment. We further show that this can be measured with increased sensitivity by sampling the evolution in the time and phase domain non-uniformly, according to a mathematically optimal adaptive experimental design. Finally, we report applications of this method to characterise dynamics within folded FLN5 RNCs.

## Results

### Analysis of cross-correlated relaxation in methyl spin systems during an F1-coupled HSQC experiment

The relaxation of methyl spin systems has been treated and reviewed on previous occasions (*23*, *26*), but for completeness in this section we give a brief summary of relevant aspects. Fig. 2 shows a diagram of the energy levels and single-quantum transitions in an isolated ^13^CH_3_-labelled methyl group. In the macromolecular limit and assuming rapid rotation about the three-fold symmetry axis, each transition relaxes in a single-exponential manner. However, the relaxation rates of individual ^1^H or ^13^C transitions vary due to cross-correlations between dipole-dipole (DD) interactions or between DD interactions and the ^13^C CSA (*26*).

There are ten ^1^H single quantum transitions present in the spin system (vertical transitions in Fig. 2). Due to cross-correlations between proton dipolar interactions (*η*_*HHHH*_), these can be divided into the slowly relaxing inner lines |2〉〈3|, |5〉〈6| and |7〉〈8|, which relax with rate 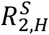, and the more rapidly relaxing outer lines |1〉〈2| and |3〉〈4|. Depending on the ^13^C spin state, these outer lines are further split by cross-correlation between HH/HC dipolar interactions (*η*_*HHHC*_) into fast (+) and slow (−) relaxing transitions, which relax with rates 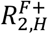 and 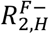 respectively (Fig. 2). Similarly, the eight ^13^C single quantum transitions (horizontal transitions in Fig. 2) relax with four different rates, 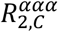, 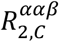, 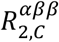 and 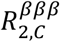, depending on the spin states of the attached protons. In particular, cross-correlations between CH dipolar interactions (*η*_*CHCH*_) lead to slow relaxation of the two inner transitions and rapid relaxation of the two outer transitions, while cross-correlation between the CH dipolar interaction and the ^13^C CSA (*η*_*CHC*_) leads to slower relaxation where attached protons are in the *β* spin state (Fig. 2). Importantly, in the macromolecular limit all of these cross-correlations depend only on the spectral density at zero frequency, *J*(0)=S^2^τ_c_, and on the ^13^C CSA, Δσ_*C*_, which depends primarily on the residue type. Values of 18.2 ± 1.5 and 25.8 ± 5.6 ppm were measured in protein L for isoleucine Cδ and leucine/valine methyls respectively (*29*). It will be demonstrated below that these mean values provide an acceptable degree of accuracy for our purposes, and therefore using this approximation all necessary CCR rates may be calculated from *J*(0) alone.

Fig. 3A shows the HSQC pulse sequence used in this work, in which the central ^1^H decoupling pulse is removed so that ^13^C multiplet structures can be resolved (Fig. 3B). The evolution of anti-phase ^13^C magnetisation during *t*_1_ would be expected to give rise to a 3:1:1:3 multiplet pattern, and deviations from this have previously been used to evaluate cross-correlated relaxation processes in lysine sidechain NH_3_ groups (*34*) and, using related constant-time evolution experiments, in methyl groups (*26*, *29*, *30*). However, we will show below that cross-correlated relaxation occurring during the remainder of the experiment may distort the expected 3:1:1:3 multiplet structure, leading to previously unappreciated systematic errors. In order to describe the amplitudes of lines within ^13^C multiplets correctly, it is therefore essential to calculate the evolution of magnetisation throughout the entire pulse sequence.

**Figure 3.**
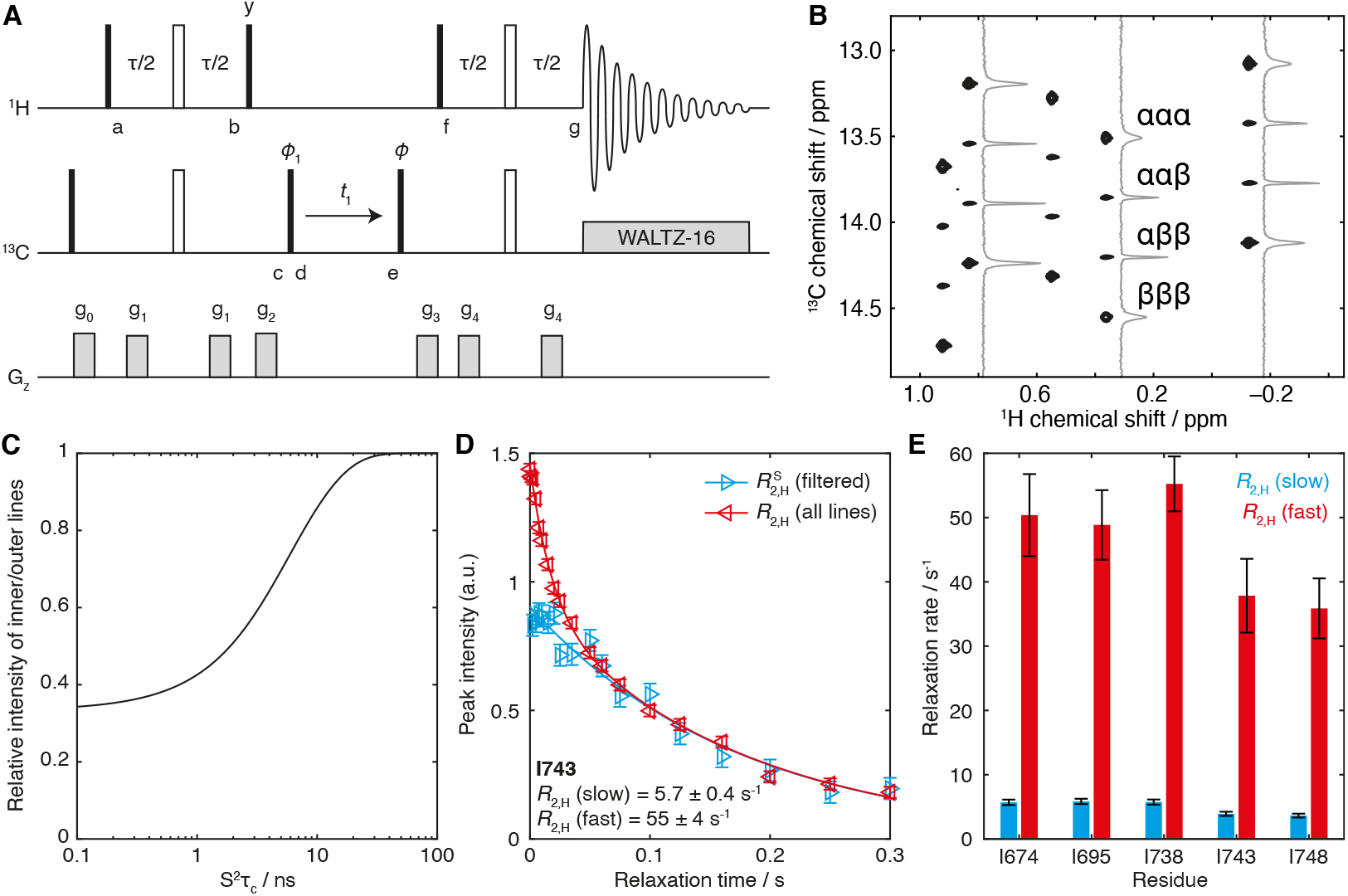
Analysis of the F1-coupled HSQC experiment. (**A**) Pulse sequence diagram of the F1-coupled ^1^H,^13^C HSQC experiment used in this work. The delay τ is 4 ms (1/2*J*_CH_), phase *ϕ*_1_ = x, −x; receiver phase *ϕ*_rx_ = x, −x. States-TPPI quadrature detection is achieved by incrementing the phase of the pulse marked *ϕ*. Alternatively, for measurements in the time domain, the phase *ϕ* can be set to any value from 0 to 180°. Gradients are applied as 1 ms smoothed squares, with strengths g_0_ = 0.0935 T m^−1^, g_1_ = 0.0385 T m^−1^, g_2_ = 0.275 T m^−1^, g_3_ = 0.19 T m^−1^ and g_4_ = 0.07 T m^−1^. (**B**) F1-coupled ^1^H,^13^C HSQC spectrum of [^2^H,^13^CH_3_-Ile]-labelled FLN5 (950 MHz ^1^H Larmor frequency) with cross-sections illustrating the differential relaxation of ^1^H spin states, as indicated, within the ^13^C multiplet. (**C**) Ratio of the initial amplitudes of inner and outer transitions, *I*_*in*_/*I*_*out*_ = (3 − Δ)/(3 + 3Δ) (Eq. 3), plotted as a function of S^2^τ_c_ with Δ calculated according to Eq. 2. (**D,E**) Measurement of slow and fast ^1^H relaxation rates for FLN5 (298 K, 700 MHz ^1^H Larmor frequency) by global fitting of ^1^H Hahn echoes including/excluding a filter for the fast-relaxing outer transitions (*37*).

Following the initial 90° H_x_ pulse, the density operator 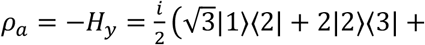 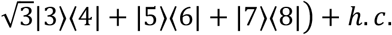 (where *h.c.* indicates the Hermitian conjugate of the expression) evolves under the scalar coupling Hamiltonian 2*πJ*_*CH*_H_*z*_C_*z*_ for a total period *τ* = 1/2*J*_*CH*_. The chemical shift is refocused during this interval, while relaxation occurs under the influence of cross-correlated HH/HH dipolar interactions and HH/HC dipolar interactions, following Fig. 2. Note that these cross-correlations are not affected by the central 180° (H_x_ + C_x_) pulse, and so at point b the density operator is 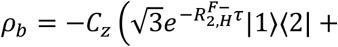 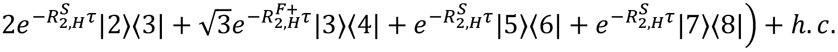.

Following the 90° H_y_ pulse, which mixes together terms in the *I* = 3/2 manifold, and the purge gradient g_2_, the density operator at point c becomes:

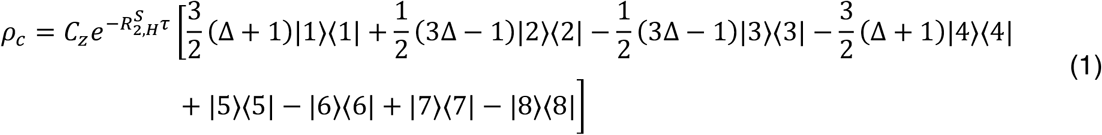

where the amplitude

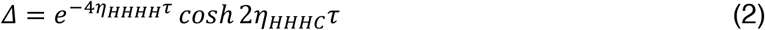

accounts for the effects of cross-correlated relaxation during the INEPT transfer. Note that Δ, like the rates *η*_*HHHH*_ and *η*_*HHHC*_, is a function of *J*(0) only, and is close to unity for small *J*(0) (i.e. rapid tumbling), decreasing towards zero as *J*(0) increases. The effect of this term, which represents longitudinal four-spin (16*C*_*z*_*H*_*z*_*H*_*z*_*H*_*z*_) order within *ρ*_*c*_, is to perturb the initial amplitudes of the inner and outer lines generated by the 90° C_x_ pulse at point d from the canonical 3:1:1:3 ratio towards 1:1:1:1 as *J*(0) increases (Fig. 3C).

From point d to e, ^13^C coherences evolve under the influence of the ^13^C chemical shift and the heteronuclear scalar coupling, with spin-state dependent relaxation rates as indicated in Fig. 2. While ^1^H spin flips induced by interactions with external spins can couple adjacent transitions, because the ^13^C chemical shift and heteronuclear scalar coupling are not refocused in this period then provided that the rate of spin flips is much less than the frequency separation between adjacent lines of the multiplet, *R*_1*H,sel*_ ≪ 2*πJ*_*CH*_, these coupling terms are non-secular and hence cross-relaxation is strongly suppressed.

The back-transfer from point f to g following the final 90° H_x_ pulse can be analysed similarly to the initial INEPT transfer. At the point of acquisition, both fast and slow-relaxing ^1^H coherences are present and will give rise to degenerate resonances in the direct dimension, with amplitudes 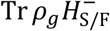, where 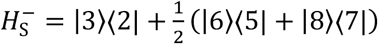 and 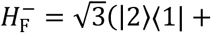 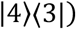. Carrying out the calculation in this manner, we find that the observed magnetisation is:

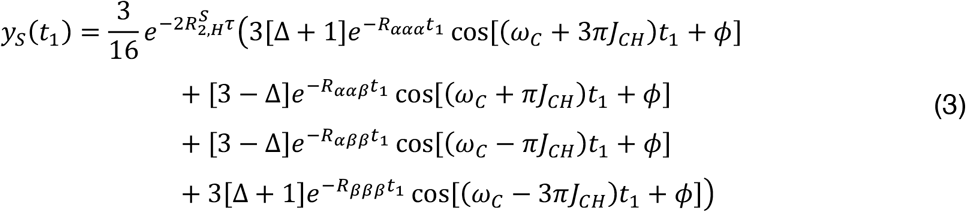

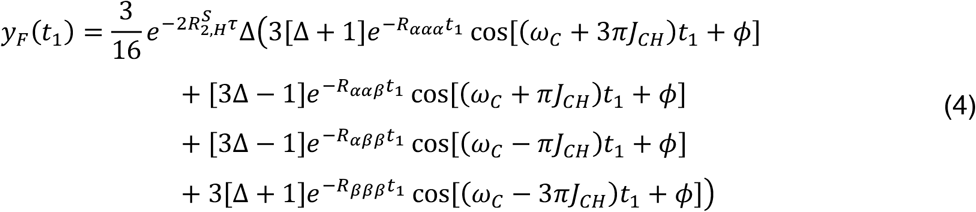

where *ϕ* indicates the phase of the final ^13^C 90° pulse, and the relaxation rates *R*_*ααα*_ etc. are defined in Fig. 2. As all signals are equally weighted by the factor 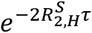, this will effectively be absorbed into an overall amplitude for each signal.

Following detection (with ^13^C decoupling) and Fourier transformation, each component above gives rise to a Lorentzian signal in the ^1^H dimension, with linewidths given by the slow and fast ^1^H relaxation rates respectively:

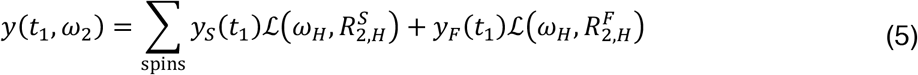

The fast-relaxing component in this expression, *y*_*F*_(*t*_1_), is suppressed by a factor Δ relative to *y*_*S*_(*t*_1_), which tends towards zero as S^2^τ_c_ increases (Eq. 2). Moreover, as the intensity of a cross-peak is inversely proportional to its linewidth, and given that 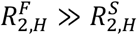 (as illustrated experimentally for measurements of isolated FLN5 in Fig. 3D,E), to an excellent approximation the fast relaxing terms may be neglected in the above expression.

Eq. 3 may be used to fit data directly in the *t*_1_ time domain, by integration around the ^1^H chemical shift of an isolated resonance. Alternatively, the Fourier transform of the expression is simply a sum of four Lorentzian signals corresponding to the four multiplet components, with amplitudes and linewidths as indicated (plus additional contributions depending on the particular window function applied), and this may be used to fit cross-sections from spectra in the frequency domain.

A final point remains to be considered regarding the effects of relaxation due to the interaction of spins with external protons, and the potential for the introduction of systematic errors. Relaxation during the evolution of ^1^H coherences (from point a to b and f to g in Fig. 3A) can be described in a Liouville subspace (for simplicity here showing only the |*α*〉 ^13^C spin state):

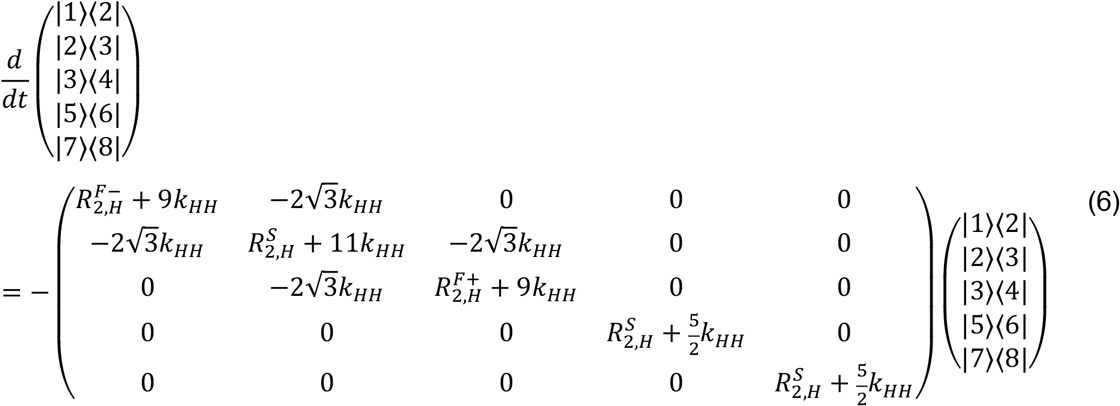

where 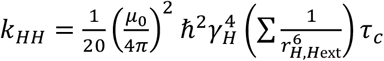 and similar relations apply for the |*β*〉 ^13^C spin state and Hermitian conjugates (*26*). In part, the additional diagonal terms can be absorbed into the definition of 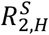, which in turn will affect only the overall amplitude of the signal. The significance of the remaining diagonal and off-diagonal terms depends primarily on the relative magnitudes of *k*_*HH*_ and 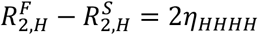. For [^2^H,^13^CH_3_-Ile], the sum over interproton distances may be approximated by the weighted average 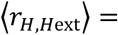 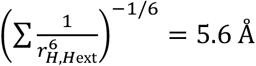, on the basis of which the relative magnitude of external proton interactions are estimated to be an order of magnitude weaker than cross-correlated relaxation processes. However, for non-deuterated proteins, in which the weighted average ̩*r*_*H,Hext*_ is considerably smaller, the effects of external spins are likely to be larger and potentially non-negligible.

Interactions with external spins also introduce an additional perturbation to the evolution of the two inner lines during *t*_1_, beyond the non-secular cross-relaxation terms already discussed. The proton density matrix elements relevant to the *ααβ* line in *ρ*_*c*_ (Eq. 1) can be re-written as 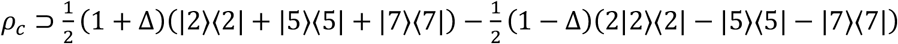 (and a similar analysis follows for the *αββ* line). When *J*(0) is small and little cross-correlated relaxation takes place during the initial INEPT transfer (Δ ≈ 1), only the first term is present. This can be expanded as |2〉〈2| + |5〉〈5| + |7〉〈7| = |*ααβ*〉〈*ααβ*| + |*αβα*〉*αβα*| + |*βαα*〉〈*βαα*|, i.e. a simple population term, which relaxes with external protons with rate 3*k*_*HH*_ (*26*). However, when *J*(0) is larger and cross-correlated relaxation during the INEPT transfer cannot be neglected, the second term may become comparable to the first. This can be expanded as 2|2〉〈2| − |5〉〈5| − |7〉〈7| = |*ααβ*〉〈*αβα*| + |*ααβ*〉〈*βαα*| + |*αβα*〉〈*ααβ*| + |*αβα*〉〈*βαα*| + |*βαα*〉〈*ααβ*| + |*βαα*〉〈*αβα*|, i.e. a symmetrised mixture of zero quantum ^1^H transitions, which relaxes more rapidly with external protons, with rate 11*k*_*HH*_ (*26*). The net effect of this is that the two inner transitions will be slightly broader than otherwise expected, leading to an underestimation of S^2^τ_c_. As above, we estimate that the impact of this effect will be small for isolated ^13^CH_3_ groups in perdeuterated proteins, but may be more significant if applied to the analysis of non-deuterated molecules. Any application to such molecules should therefore be carried out with attention to these systematic effects.

### Experimental validation

Prior to application to RNCs, we carried out a series of measurements using isolated [^2^H,^13^CH_3_-Ile]-labelled FLN5 (Fig. 1B) to validate the accuracy of our analysis, against reference measurements acquired using the triple quantum build-up scheme (*28*). Measurements were performed in a series of d_8_-glycerol concentrations, at 298 K and 283 K, in order to decrease the rate of tumbling and so probe rotational diffusion across a wide range of timescales. Fig. 4A–D shows coupled HSQC spectra acquired under these conditions, together with HMQC reference spectra for the determination of peak positions.

**Figure 4.**
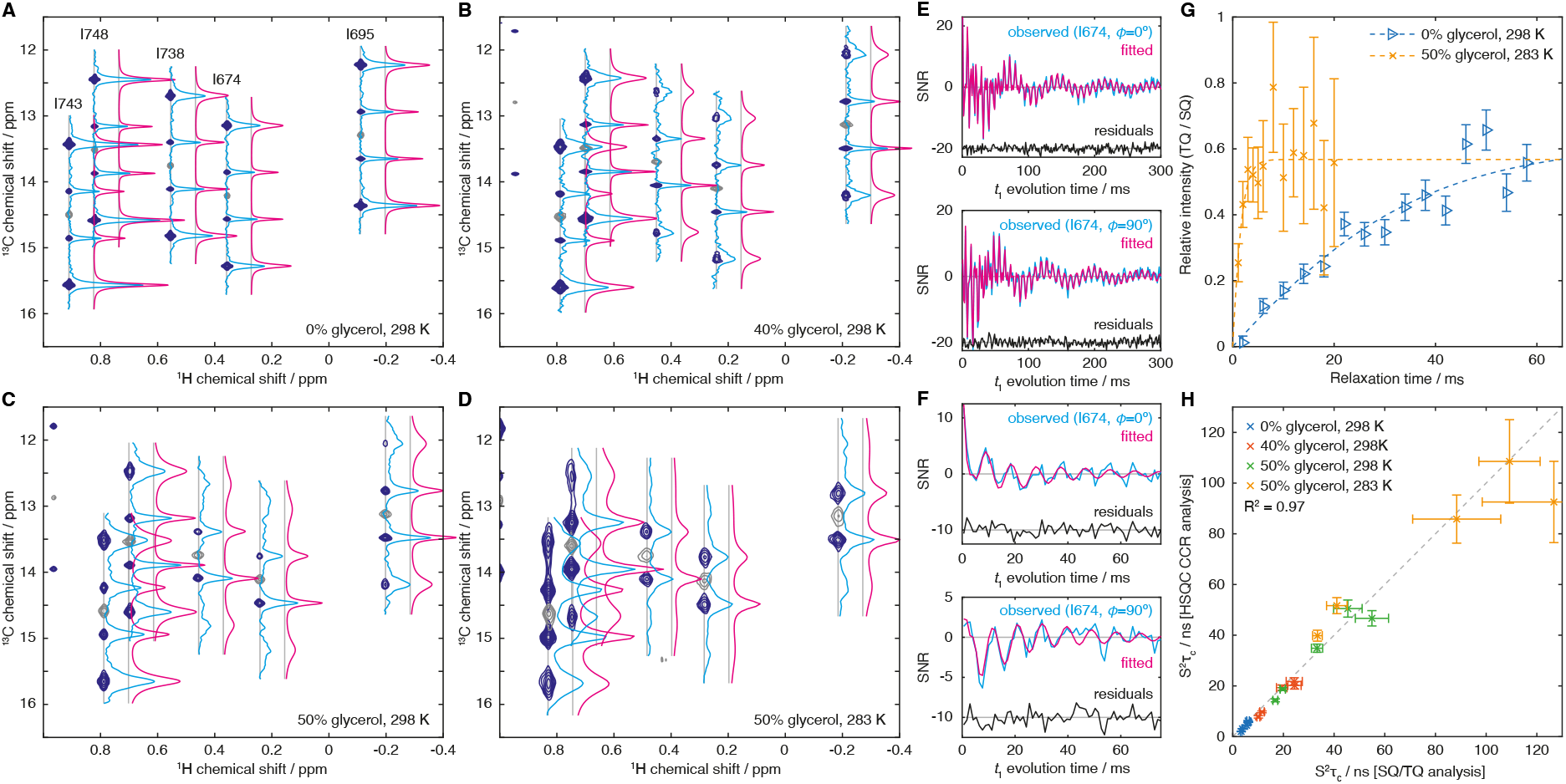
Experimental measurements of methyl group dynamics in isolated FLN5. (**A–D**) F1-coupled ^1^H,^13^C HSQC spectra (Fig. 3A, dark blue) and reference ^1^H,^13^C HMQC (grey) acquired for isolated [^2^H,^13^CH_3_-Ile]-labelled FLN5 at varying temperatures and d_8_-glycerol concentrations as indicated. Spectra were processed with exponential window functions in the indirect dimension (5 Hz (A), 5 Hz (B), 10 Hz (C) and 20 Hz (D)). Cross-sections through ^13^C multiplets are shown with blue lines, and fits to the Fourier transform of Eq. X, incorporating the effects of the window function, are shown in magenta (horizontally offset for clarity). (**E–F**) Time domain analysis of relaxation measurements, showing the observed *t*_1_ evolution of cosine (*ϕ*=0°) and sine (*ϕ*=90°) modulated components of the I674 resonance, in (E) 0% glycerol, 298 K, and (F) 50% glycerol, 283 K (blue lines). Cosine and sine modulated components were fitted simultaneously to Eq. 3 (magenta lines). (**G**) Reference measurements of S^2^τ_c_ in isolated FLN5 by measurement of the build-up of triple quantum magnetisation (*28*). Data and fits are shown for the I674 resonance in 0% glycerol, 298 K, and 50% glycerol, 283 K, as indicated. (**H**) Comparison of S^2^τ_c_ measurements determined by both experimental approaches. Data are shown for all five isoleucine groups of FLN5, with solution conditions as indicated.

The spectrum of FLN5 in the absence of glycerol (298 K, Fig. 4A) shows strong outer lines, although it can be observed that these are broader than the less intense inner lines. As the glycerol concentration is increased, these outer lines become broader and less intense, and the asymmetry under exchange *α* ↔ *β* introduced by cross-correlation with the ^13^C CSA starts to become discernible (Fig. 4B,C). At the highest glycerol concentration and lowest temperature acquired (Fig. 4D), for several residues the outer lines are barely observable, but the asymmetry between the inner *ααβ* and *αββ* lines nevertheless may be expected to provide a continued route to the quantification of rotational diffusion.

The extracted cross-sections were fitted to the Fourier transform of Eq. 3 and are plotted in Fig. 4A–D. Equivalent results could also be obtained by direct fitting of time-domain data (Fig. 4E–F). The ^13^C chemical shift was determined from the peak position in the accompanying HMQC spectrum, the ^13^C CSA was fixed to 18.2 ppm as previously reported for isoleucine Cδ spins (*29*), and the heteronuclear scalar coupling *J*_*HC*_ was fixed to 125 Hz. Therefore, each cross-section could be fitted as a function of only three parameters: an amplitude factor, the ^13^C *R*_2_ relaxation rate, and S^2^τ_c_, from which all necessary cross-correlated relaxation rates could be calculated. S^2^τ_c_ values were also determined by measurement of the build-up of triple quantum magnetisation relative to the relaxation of single quantum magnetisation (Fig. 4G) (*28*). A comparison of the two measurements shows excellent agreement across two orders of magnitude in S^2^τ_c_ (Fig. 4H), thus validating further applications of the new approach.

### Optimal design of experimental measurements

In the previous sections, we have established and validated the measurement of cross-correlated relaxation in methyl groups by analysis of the differential relaxation of transitions in the ^13^C multiplet, and we have also shown that this analysis may be carried out in both the frequency (Fig. 4A–D) and time domains (Fig. 4E–F). From this point onwards, we will consider the analysis of time-domain data only, obtained following integration of non-overlapping resonances in the ^1^H frequency dimension. As such data will be fitted directly, without Fourier transformation, there is no longer a requirement either to sample uniformly in the time domain or to acquire cosine- and sine-modulated pairs of points for quadrature detection. Instead, measurements may be acquired with arbitrary *t*_1_ evolution times and phases *ϕ*, and so it is natural to ask whether a non-uniform sampling strategy might exist that provides greater sensitivity than conventional uniform sampling. Such a problem can be solved using the theory of optimal experimental design (*38*).

As we are in general considering the analysis of multiple resonances, we write our earlier expression for the time-evolution of observed signals (Eq. 3) as the vector-valued function ***y*****(*x*; *θ*)**, termed the ‘response surface’, which describes the observed intensities of *k* spins as a function of the independent variables ***x*** = (*t*_1_, *ϕ*) and the *m* = 3*k* unknown parameters 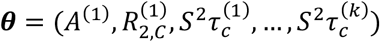. Therefore, our goal is to determine the optimal set of such experiments to perform, within the allowed space of experiments, 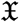, in order to estimate the parameters ***θ*** with minimal uncertainty.

We begin by defining an *N*-point experimental design *ξ*_*N*_ as a distribution of *N* measurements across *n* points 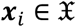, with *r*_*r*_ repeated measurements at each point *i* (∑_*i*_ *r*_*i*_ = *N*):

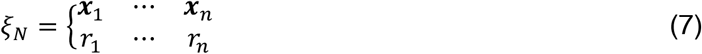

Having made a set of observations ***y***_*i*_ according to the experiment *ξ*_*N*_, where the (uncorrelated) measurements have covariance matrix σ^2^***I***_*k*_, the data can be fitted to the response surface in a least-squares sense to yield the maximum likelihood estimator of the parameters, 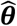. The Fisher information matrix

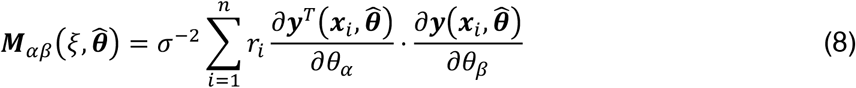

describes the curvature of the log-likelihood function about 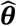, from which the parameter covariance matrix 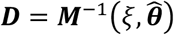 may be defined. Note that the inverse of the Fisher information matrix is the Cramér-Rao lower bound for the variance of an unbiased estimator (understood as a comparison of positive definite matrices) and so indicates the maximum achievable precision for a given experimental design. This is also independent of the noise level, which simply scales the overall uncertainty.

To determine the *prediction* error for the response surface based on this estimate (required below), we consider a linearisation of the response surface about 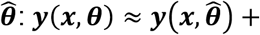 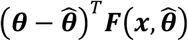, where *F*_*αβ*_ = ∂*y*_*α*_/∂*θ*_*β*_ is the Jacobian of the response surface. As the errors in the parameter estimates are normally distributed with covariance ***D***, from the transformation properties of multivariate normal distributions the predicted value of the response surface at point ***x*** is also normally distributed, 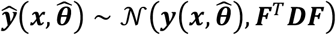. We thus define the prediction uncertainty 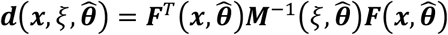.

To proceed further, we must now make our idea of the ‘optimality’ of an experiment mathematically precise. An intuitive and useful definition is to find the design *ξ*\* which minimises the volume of the parameter confidence ellipsoid, det ***D***:

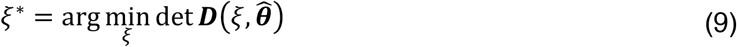

This strategy is termed *D-optimal* design and is particularly attractive as it is invariant to linear changes of parameterisation and units. It can be proved (the General Equivalence Theorem (*39*)) that this problem is also equivalent to minimisation of the maximum prediction error (termed *G-optimality*):

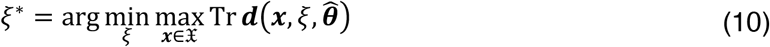

This provides an intuitive design strategy: acquire measurements at points where the prediction uncertainty is greatest, and this in turn will minimise the uncertainty in the parameter estimates. However, there is a difficulty in this approach that results from the dependence of 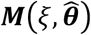 on the unknown or uncertain parameter estimate 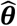, which introduces a circularity into the design. This can be resolved using an *adaptive* design (also referred to as a sequential design (*38*)) in which, following a set of seed measurements *ξ*_*N*_, an updated parameter estimate is calculated and used to select the next measurement point:

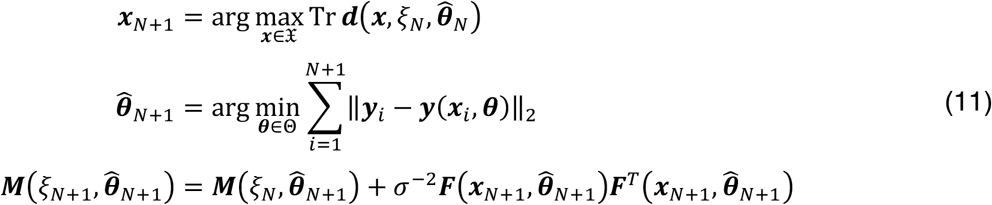

This process is illustrated as a flowchart in Fig. 5A.

**Figure 5.**
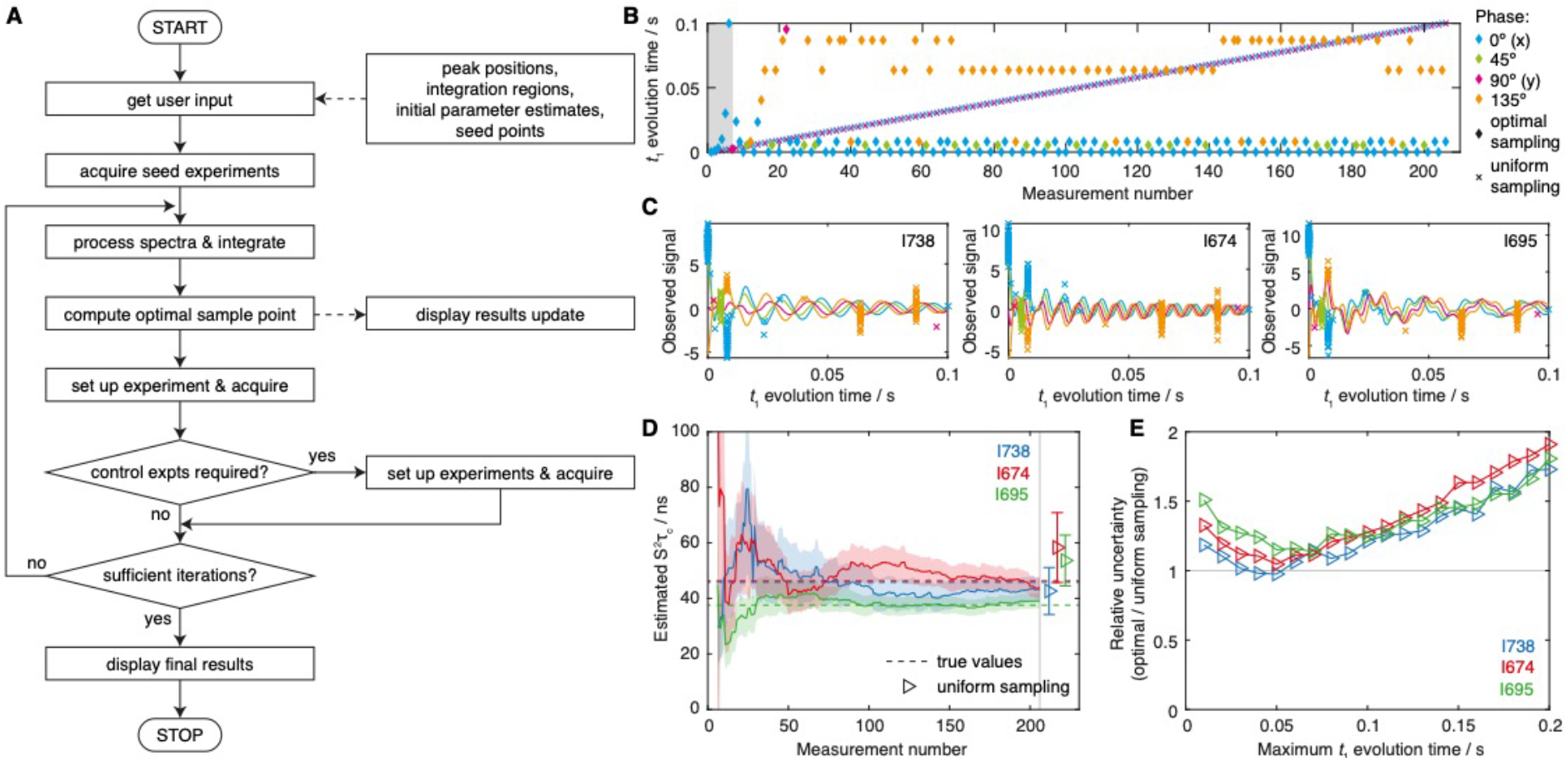
Adaptive experimental design strategy and validation using synthetic data. (**A**) Flowchart illustrating the adaptive sampling acquisition process. (**B**) Measurement points acquired during a simulated experiment, following an optimal adaptive sampling scheme or using conventional uniform sampling. The acquisition of seed measurements is marked with grey shading, while the measurement phase, *ϕ*, is shown with colour coding as indicated. (**C**) The final set of observed measurements (normalised by the noise level) are shown fitted to Eq. 3, with the measurement phase, *ϕ*, coloured as in panel B. (**D**) Evolution of S^2^τ_c_ parameter estimates as a function of the number of measurements (solid lines). Shading is used to indicate the experimental uncertainty (± s.e.). The true values used in generating the synthetic data are indicated with dashed lines, while estimates obtained from a matched, uniformly sampled dataset are shown as points with error bars (horizontally offset for clarity). (**E**) Precision of S^2^τ_c_ parameter estimates obtained using a uniform sampling scheme, with maximum *t*_1_ evolution time as indicated, relative to those determined using an optimally designed adaptive sampling scheme. Uncertainties were determined as the mean standard error across 500 independent simulations. The total number of measurements was identical for all experiments.

We initially tested the adaptive experimental design strategy using simulations to generate synthetic data. One such example is illustrated in Fig. 4B–D, using the uniformly sampled measurements of FLN5 relaxation rates obtained for I738, I674 and I695 in 50% d8-glycerol, 298 K (Fig. 3C) as the unknown ‘true’ values used to generate simulated samples. Six seed points were used to determine initial parameter estimates, with evolution times of 0.1, 1, 3, 10, 30 and 100 ms, and zero phase, following which 200 further samples were acquired according to the adaptive sampling scheme outlined above (Fig. 5A, B). The prediction error 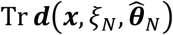 was found to be a very rough surface, and so to ensure the global maximum was located a grid search was employed across *t*_1_ values from 0 to 100 ms and phases *ϕ* ∈ {0°, 45°, 90°, 135°}. For comparison, a uniformly sampled experiment was also simulated, with an identical number of total samples and a maximum *t*_1_ acquisition time of 100 ms (Fig. 5B).

The final set of simulated measurements and the fitted evolution curves are shown in Fig. 5C, while the evolution of the parameter estimates is plotted against the increasing number of measurements in Fig. 5D. From this, the progressive reduction in the parameter uncertainty may be observed, as well as the convergence of parameter estimates towards their ‘true’ values. For comparison, estimates determined by the equivalent uniformly sampled measurement are also plotted and may be seen to have greatly reduced precision.

To explore this in more detail, 500 independent simulations of the adaptive sampling scheme were carried out in order to obtain reliable estimates of the parameter uncertainty. A series of uniformly sampled measurements were then also simulated with acquisition times ranging from 10 to 200 ms (and a fixed total number of measurement points). The relative uncertainty in S^2^τ_c_ estimates is plotted for each residue as a function of the acquisition time in Fig. 5E. It may be observed from this that the performance of uniformly sampled measurements never exceeds that of the adaptive sampling scheme. Where the optimal acquisition time is employed, the gains in precision are modest, but in general this optimal time may be different for different residues – and is of course unlikely to be known prior to carrying out the measurement. Thus, the application of adaptive sampling schemes is likely to provide a modest but useful increase in experimental precision for a fixed total measurement time.

### Measurement of cross-correlated relaxation in a FLN5+67 ribosome–nascent chain complex

Having established and validated the new experimental analysis and adaptive sampling scheme, we applied it to measure the rotational diffusion of the folded FLN5 domain within the FLN5+67 RNC (Fig. 1A). In the case of FLN5 RNCs, four isoleucine resonances can be observed and distinguished from ribosome background signals in 2D NMR experiments, and three can still be resolved unambiguously in 1D spectra (Fig. 1C,D). While this may not be a large number of resonances, because folded domains tumble as rigid bodies even a small number of observations are sufficient to provide useful insight into their motion (in contrast to disordered states, the mobility of which may vary substantially along the sequence).

Adaptively sampled measurements of I738, I674 and I695 resonances within an FLN5+67 RNC sample were acquired using six seed points to provide initial parameter estimates, with *t*_1_ evolution times of 0.02, 1, 2, 5, 10 and 100 ms. Subsequent measurements were then interleaved with control measurements of translational diffusion, and together with western blotting analysis of a sample incubated in parallel with NMR measurements these were used to infer a sample lifetime of ca. 40 hours, corresponding to 120 adaptively-sampled measurements (Fig. 6A–C). Examination of estimated S^2^τ_c_ values as a function of the number of measurements indicates that the measured values do not depend strongly on the choice of this cut-off (Fig. 6D).

**Figure 6.**
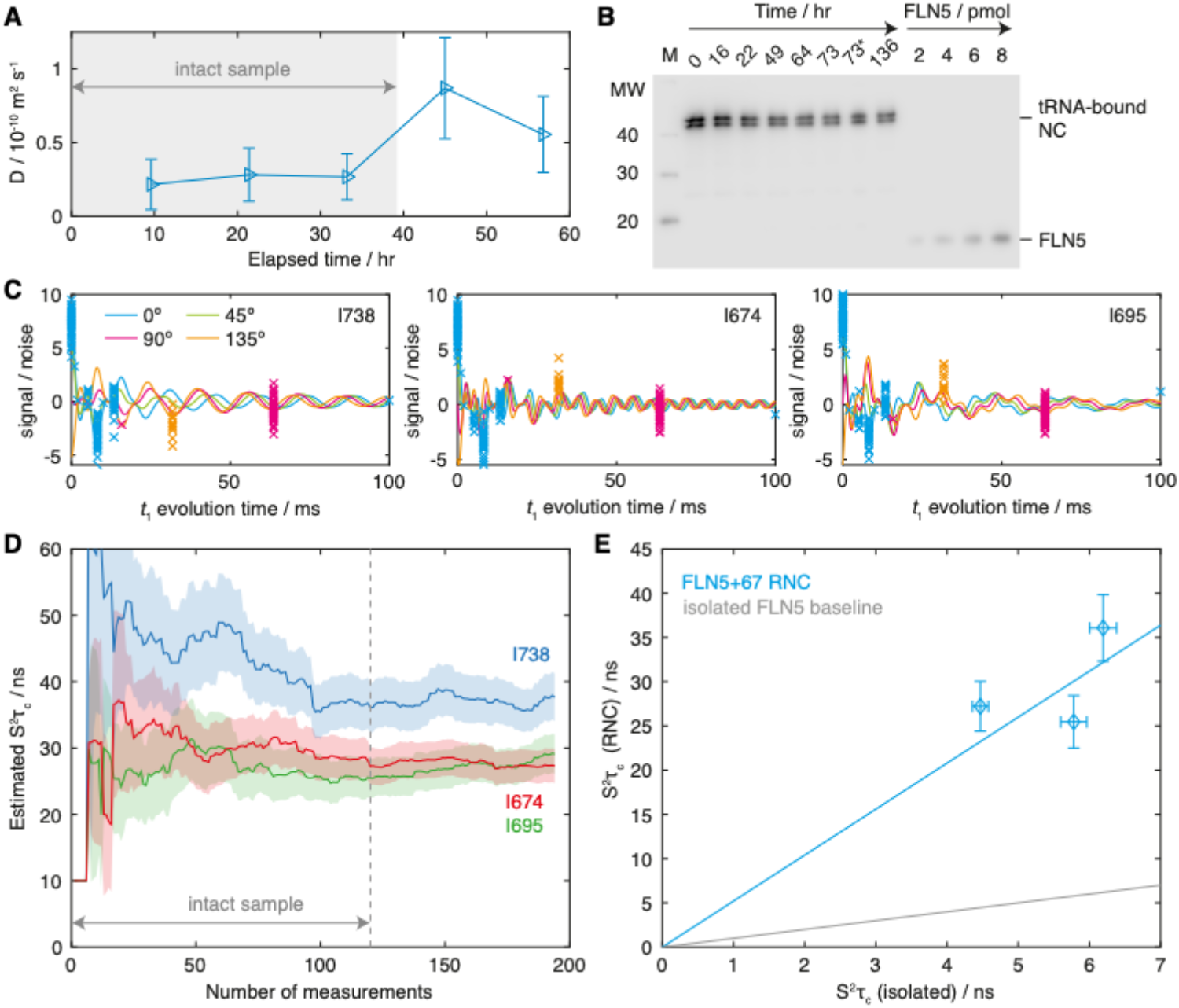
Measurement and analysis of methyl S^2^τ_c_ values in the FLN5+67 RNC. (**A**) Time course of NC diffusion coefficients measured for the isolated I738, I674 and I695 resonances. Grey shading indicates the total measurement time selected for further analysis. (**B**) Western blot (anti-histidine) time course of the FLN5+67 RNC, showing aliquots taken periodically from a sample of the RNC incubated in parallel with NMR measurements. The asterisk indicates an aliquot taken from the actual NMR sample. The locations of the higher molecular weight tRNA-bound NC, and of isolated FLN5, are indicated. (**C**) Adaptively sampled F1-coupled HSQC measurements for I738, I674 and I695 resonances of the FLN5+67 RNC (normalised by the noise level) are shown fitted to Eq. 3, with the measurement phase, *ϕ*, coloured as indicated. (**D**) Evolution of S^2^τ_c_ parameter estimates as a function of the number of measurements (solid lines). Shading is used to indicate the experimental uncertainty (± s.e.). The dashed line indicates the cut-off for sample integrity, as determined by diffusion and western blot analysis (panels A and B). (**E**) Comparison of isoleucine S^2^τ_c_ values measured for the FLN5+67 RNC and for isolated FLN5. A regression line through the origin is shown in blue (with a gradient of 5.2), and a reference line for the isolated domain is shown in grey.

The S^2^τ_c_ values determined from this measurement were 37.7 ± 3.6 ns (I738), 29.1 ± 3.0 ns (I674) and 27.3 ± 2.6 ns (I695). By comparison, S^2^τ_c_ values of 6.2 ± 0.2 ns (I738), 5.8 ± 0.2 ns (I674) and 4.5 ± 0.1 ns (I695) were determined for the isolated domain under equivalent conditions. The values measured for the NC are instead approximately intermediate between measurements of isolated FLN5 in 40% and 50% glycerol. Assuming that S^2^ order parameters are not altered within the NC – which seems unlikely given that no chemical shift perturbations are observed – then these differences may be interpreted in terms of relative changes in the rotational correlation time alone, and a regression analysis indicates that rotational diffusion of the NC is approximately 5.2 times slower than the isolated domain (Fig. 6E). While the correlation between RNC and isolated measurements is not perfect, given the experimental uncertainties that we have determined together with the small number of methyl groups observed, at this stage there does not appear to be strong evidence of site-specific effects that would indicate changes in the shape of the diffusion tensor as well as its magnitude.

## Discussion

In this manuscript, we have presented a careful analysis of a previously described approach for the analysis of rotational diffusion in isolated methyl and other AX_3_ spin systems based on the fitting of multiplet lineshapes or intensities in coupled HSQC or constant-time HSQC experiments (*26*, *29*, *30*, *34*). This has revealed previously unappreciated subtleties associated with the effects of cross-correlation relaxation during the entirety of the sequence, resulting in initial multiplet amplitudes that deviate from a simple 3:1:1:3 ratio for slowly tumbling systems (Fig. 3C). By incorporating this correction, we have shown that values of S^2^τ_c_ may be determined accurately over a wide range of correlation times, from 1 ns to over 100 ns (Fig. 4H).

We have next developed and implemented an on-the-fly adaptive sampling scheme for this measurement based on the principles of optimal experimental design (*38*). The optimal design of NMR experiments has been discussed in many previous works, generally in terms of the Cramér-Rao lower bound (the inverse of the Fisher information matrix ***M***(*ξ*)), although the precise criteria employed to optimise this matrix quantity are not always clearly stated. An early application to protein NMR spectroscopy was the optimal selection of time points for relaxation measurements (*40*), while applications of the CRLB to parametric estimation of multidimensional NMR spectra, and the evaluation of non-uniform sampling schemes, was reported a few years later (*41*). More recently, this approach has been applied to droplet size estimation using diffusion NMR experiments (*42*), and to the optimal parametric estimation of protein relaxation rates in a non-uniformly sampled accordion NMR experiment (*43*). However, to the best of our knowledge the only previous report of an adaptive approach to design and acquire new NMR experiments ‘on-the-fly’ is ADAPT-NMR, used for automated protein resonance assignment (*44*). The combination of the two approaches that we present here has demonstrated the increased precision or reduced acquisition time that can be provided by these methods (Fig. 5E), and points towards potential to applications of adaptive optimal designs to other areas of NMR such as relaxation measurements or non-uniform sampling.

We have applied the methods developed to the analysis of rotational diffusion within a ribosome–nascent chain complex. As noted in the introduction, understanding the dynamic behaviour of NCs as they emerge from the ribosome during biosynthesis is a key challenge in order to elucidate the mechanisms by which the ribosome may modulate or regulate the co-translational folding process (*4*). The slower tumbling of the FLN5+67 NC that we have observed may reflect contributions from a number of factors. Firstly, the presence of the FLN6 linker may reduce the mobility of the attached domain. We have previously noted that amide resonances of this linker are strongly broadened in RNC spectra across many lengths (*9*), which may indicate the existence of interactions between the linker and the ribosome surface, or a conformational exchange process such as transient or partial folding. Such processes might in turn be expected to hinder the rotational diffusion of the attached domain. Moreover, if the FLN5 domain is unable to sample all orientations equally due to restrictions from the linker, then the *J*(0) spectral density will have an additional contribution reflecting this restricted tumbling, proportional to the ca. 2.5 *μ*s rotational correlation time of the parent ribosome (*12*).

Lastly, the slower tumbling we observe may indicate transient interactions between FLN5 and the ribosome surface. Assuming rapid exchange, the observed rotational correlation times will be a population weighted average of those in the free and bound states, and from this we may infer that the maximum population of any bound state is approximately 1.5% (and additional contributions to the slower tumbling as discussed above would limit this population further). In this instance, such a weak interaction with the largely negatively-charged ribosome surface (*45*) is perhaps understandable given that the total charge on the FLN5 domain is –9. Indeed, using both NMR and fluorescence methods, ribosome–NC interactions have previously been identified to be driven, at least in part, by electrostatic interactions (*7*, *10*). Nevertheless, a detailed understanding of such weak interactions, as well as other dynamic effects, may be expected to yield fundamental insights into the behaviour of folded NCs as they emerge from the ribosome, and the optimised experimental approaches that we have developed here will facilitate a new generation of sensitive and precise studies of these phenomena.

## Materials and Methods

### Sample preparation

Isolated [^2^H,^13^CH_3_-Ile]-labelled FLN5 was expressed and purified using standard methods, as previously described (*9*). The FLN5+67 RNC construct (Fig. 1A) has also been previously described, and was expressed using minor adaptations of previously reported methods (*9*, *17*). *E. coli* BL21 (DE3) cells were progressively adapted to EM9 medium containing 100% D_2_O and d_7_-glucose, and then grown to an OD_600_ of 3. Selective isoleucine methyl labelling of the nascent chain was achieved by addition of 80 mg/L of the isoleucine precursor 2-ketobutyric-4-^13^C,3,3-d_2_ acid 30 minutes before induction with 1mM IPTG for 50 min at 310 K. RNCs were purified from harvested cells following our previously reported protocols, except for the substitution of a second sucrose cushion in place of the final sucrose gradient purification step (*9*). Finally, RNCs were flash frozen and stored at −80°C prior to NMR measurement.

### NMR measurements of isolated FLN5

Three samples of [^2^H,^13^CH_3_-Ile]-labelled FLN5 (ca. 100 *μ*M) were prepared in deuterated Tico buffer (100% D_2_O and d_8_-HEPES, pH 7.5) containing 0, 40 and 52% (w/w) d_8_-glycerol. NMR experiments were acquired at using a Bruker Avance III HD spectrometer operating at a ^1^H Larmor frequency of 700 MHz. ^1^H,^13^C HMQC and (uniformly sampled) F1-coupled ^1^H,^13^C HSQC experiments (Fig. 3A) were acquired with acquisition times of 107 ms in the ^1^H dimension and spectral widths of 20 ppm and 2.4 ppm in the ^1^H and ^13^C dimensions respectively. 384, 256, 256 and 64 complex points were acquired for 0%, 40%, 52% and 52% (283 K) samples respectively. Measurements of triple quantum build-up and single quantum relaxation were acquired for each sample and fitted as previously described (*28*). All experiments were processed using nmrPipe (*46*) and analysed using MATLAB.

### Implementation of adaptive sampling

Adaptive sampling methods were implemented for Bruker spectrometers operating Topspin 3.5pl6. While a Python interface is provided within Topspin, this runs using the Java implementation, Jython, which does not support the required numerical libraries. Therefore, a separate instance of Python 3 is required, together with installations of NumPy, SciPy and Numba (*47–49*). Simple routines have been developed to interface between Topspin and Python, using json formatted files to exchange data. All necessary code, scripts and pulse sequences are available online (https://github.com/chriswaudby/adaptive-sampling).

### NMR measurement of the FLN5+67 ribosome–nascent chain complex

Immediately prior to NMR measurement, RNC samples were buffer exchanged into deuterated Tico buffer (100% D_2_O and d_8_-HEPES, pH 7.5) supplemented with protease inhibitors (Roche), with a final sample concentration of 7.5 μM. Some of the sample was reserved in order to monitor the integrity of the NC over the course of the NMR measurements. This control sample was incubated at 298 K, and aliquots containing 2 pmol RNC were periodically flash frozen until NMR data collection was complete. These samples were analysed by anti-His western blotting, and the intensity of the band corresponding to tRNA-bound NC was determined relative to isolated standards using densitometry (with ImageJ software, http://imagej.nih.gov/ij/).

RNC NMR experiments were performed at 298 K using a Bruker Avance III HD spectrometer running Topspin 3.5pl6 and operating at a ^1^H Larmor frequency of 950 MHz. F1-^1^H-coupled HSQC measurements were acquired with adaptive sampling as described above, with 2048 complex points in the direct dimension (107 ms acquisition time), 512 scans per measurement, and an inter-scan delay of 1.5 s (total measurement time 15 min). RNC integrity was monitored during data collection using ^13^C-filtered ^1^H STE diffusion measurements (*50*), which were interleaved between every six HSQC measurements, using a diffusion delay of 50 ms, bipolar smoothed-square gradient pulses with total length 4 ms and strengths of 0.024 and 0.457 T m^−1^, 512 scans and an inter-scan delay of 1.5 s (total measurement time ca. 30 min). The combined integrals of the isolated RNC methyls I738, I674 and I695 were analysed to provide a specific probe of the attachment of the NC.

## Acknowledgements

We acknowledge the use of the UCL Biomolecular NMR Centre and the staff for their support. This work was supported by the Francis Crick Institute through provision of access to the MRC Biomedical NMR Centre. The Francis Crick Institute receives its core funding from Cancer Research UK (FC001029), the UK Medical Research Council (FC001029), and the Wellcome Trust (FC001029). This work was supported by a Wellcome Trust Investigator Award (to J.C., 206409/Z/17/Z) and the BBSRC (BB/T002603/1). This study made use of NMRbox: National Center for Biomolecular NMR Data Processing and Analysis, a Biomedical Technology Research Resource (BTRR), which is supported by NIH grant P41GM111135 (NIGMS) (*51*).

